# Assessment of diseases and insect pests of apple (*Malus domestica* Borkh) in Silte Zone, Southern Ethiopia

**DOI:** 10.1101/2023.06.22.546144

**Authors:** Sebsibe Sh. Woto, Adimasu A. Aloto

## Abstract

Assessment research was conducted in six *Kebeles* (administrative units) of Mirab azerenet and Alicho wuriro Districts of Silte Zone in 2021/21 cropping season, to assess the prevalence and damage of diseases and insect pests of apple and growers experience on pest identification and its management. A total of 60 apple growers were interviewed using a semi-structured questionnaire. Powdery mildew and apple scab were more damaging diseases of apple fruit tree in the area and they imply economic threat if not managed and also green apple aphid was the only recorded insect pest that caused leaf curling of apple fruit tree. About 71.7% of respondents were responded powdery mildew as number one disease of apple that hinders apple production. About 76.7% of producers were answered that they didn’t identified insect pests that attack their apple fruit tree. Out of the total respondents 90% (for diseases) and 16.7% (for insects) responded that, winter season was primary period for occurrence of these pests in their apple orchard/nursery. Out of the respondents 61.7% and 85% (for diseases and insect pests) responded that they don’t have any information about management methods. About 85% of respondents replied that, they don’t prune their apple fruit tree in the optimum season and also they don’t know its importance and the method. Generally, the result of current assessment revealed that, awareness of growers on crop agronomy, pest type and its infestation season, and management measures that taken so far were not encouraging. Based on the results of our survey, we recommend the following; timely awareness of apple fruit tree; agronomy, improvement and protection should be provided for growers by agricultural extension workers and other agents.

## Introduction

Apple (*Malus domestica* Borkh) is a woody plant belonging to the family, Rosaceae and subfamily Maloideae or formerly Pomoideae [1; cited by 2]. It is the world’s deciduous fruit which accounts for 50% of fruit tree production globally putting China at the front of the world’s apple producing country; followed by USA, India, and Turkey. The leading producer of Apple in Africa is South Africa followed by Egypt and Kenya [3, 4]. In Ethiopia, apple was first brought to Southern Ethiopia, Gamo Zone, Chencha District by Missionaries before 60 years ago and now a day it grown in different highland parts of the country [2].

In Ethiopia, production of horticultural crops is much less developed and also management knowledge and skills demanded by the apple plant is not yet well familiarized among experts [4]. Total production of apple at national and regional level is not reported on [5] data and similar idea is raised by [2], there is no statistical evidence on the total areas covered and annual production of temperate fruits in Ethiopia. In Silte Zone farmers produce over 16,000 quintals of apple per year (specifically; Mirab Azernet Berber District produce 300 quintals of apple per year and Alicho Wuriro District produce 113.5 quintals of apple per year). Out of the Zone’s 480 hectares of land that are suitable for apple production, only 54 hectares have so far been cultivated [6,7].

Production and sales of apple fruits have great contribution to the economy of farmers in Ethiopia and nowadays, the cultivation of apple trees by farmers is increased because of its importance as a source of income, since apple fruits as well as apple rootstocks used for grafting of scions are propagated and sold for apple growers within the country in relatively high price [8]. Producing apple is highly promising and financially feasible both in terms of fruits and seedlings production, and becoming an interesting business for both rural and urban smallholders [9, 10].

The report of different researchers on pests that attack temperate fruits in Ethiopia shows that, apple fruit trees are subjected to different insect pests (like; aphids, caterpillars, plant bugs, and maggots) and diseases (like; powdery mildew, apple scab, leaf rust, canker, and root rot) which attack its leaf, flower, fruit, stem and root [11,10]. Even though, apple fruit tree is grown in Silte Zone, there is a lack of information on apple production limiting factors especially on diseases and insect pest’s infestation, damage and farmers’ experience on insect pest and disease identification and management. Therefore, the current research was planned to deal with the objective, to assess prevalence and damage of diseases and insect pests of apple and growers experience on pest identification and its management.

## Materials and methods

### Description of the study area

The survey was conducted in Silte Zone, Southern Ethiopia in two rural Districts namely; Mierab azernet berber and Alicho wuriro and Districts were selected due to their apple production potential. They were geographically located at 7° 43’ 30’’ North and 37° 54’ 44’’ East 38° and 7° 56’ 416’’ North and 09’ 048’’ East. The altitude ranges up to 2601-3100 and 2700-3000 [12].

### Field survey and sampling

From two Districts, in the first stage major pome/apple fruit producing *Kebeles* were selected purposively in discussion with the respective District agricultural offices. In the second stage, three model apple producing *Kebeles* were selected randomly from each District and a total of six apple producing *Kebeles* were selected from two Districts and also ten apple orchards per *Kebeles* were selected randomly with total of thirty farmers apple orchard per District. A total of 60 farmer apple orchards were used for the survey.

Similarly, a lottery method procedure was adopted for selection of three apple trees per farmer field as per [13] cited by [14] and 30 apple trees per *Kebele* was selected and a total of 180 apple trees were assessed. For the assessment of diseases and insect pests, shoots and fruits infested with different pests were counted and per tree percent infestation was computed [15]. Each selected farmers apple orchards were surveyed in two week interval (two times a month). Pests were identified in the field using identification keys (visual observation of signs and symptoms of diseases and insect pests, using a pocket/hand lens and guide line books/literatures).

### Data collected

Prevalence of diseases and insect pests in farmer’s apple orchard were recorded by the research team. The insect damage scale was noted according to [16] that states very low (≤5%), 1ow (6 to 10%), medium (11 to 20%), high (21 to 50%) and very high (>50%) levels. Infestation and damage level from attacked plant parts were calculated by using the following formula.

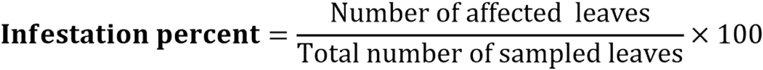

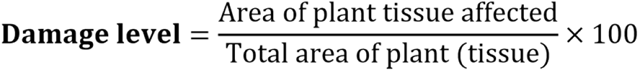

Disease severity was recorded by using 0-5 visual diseases scoring scale stated by [17]. Where; 0 = no symptoms (healthy), 1 = slight, less than 5% of leaves affected, 2 = moderate, 5-20% of leaves affected, some yellowing, little or no defoliation, 3 = extensive, 20-50% of leaves affected, significant defoliation, 4 = heavy, 50-80% of leaves affected, severe defoliation and 5 = extreme, 80-100% leaves affected, complete or near complete defoliation.

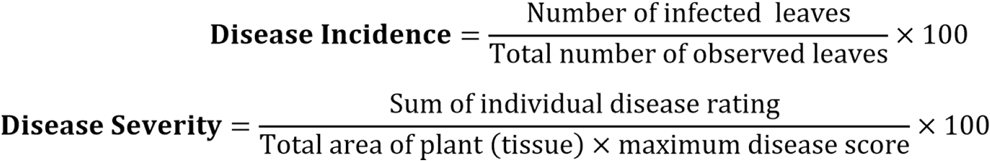

Damage/severity of fruits were rated by 0-3 scale and average percentage respectively for each were; 0 = no symptom (healthy), 1 = slight (less than 5% of fruits affected), 2 = moderate (5-20% of fruits affected, some yellowing, little or no defoliation), 3 = extensive damage (20-50% of fruits affected) [18]. Secondly, a semi-structured questionnaire was developed to collect data on apple tree management and awareness of farmers on occurrence of diseases and insect pests and their control methods. Respondents were interviewed by trained enumerators with supervision of the research team.

### Method of data analysis

The collected data was coded and entered in SPSS software version 20. Descriptive statistics were computed and the output presented in the form of percentages using tables.

## Results

### Diseases and insect pests incidence and severity level

Different diseases and insect pests of apple were identified that affecting apple fruit tree in Silte Zone, Southern Ethiopia. Among diseases recorded were powdery mildew (*Podosphaera leucotricha*), apple scab (*Venturia inaequalis*), fire blight (*Erwinia amylovora*) from leaf of apple and again powdery mildew and apple scab were recorded from fruit of apple and in addition to these, green apple aphid (*Aphis pomi*) was recorded from leaf part of apple in the study area (Fig 1).

**Fig 1.**
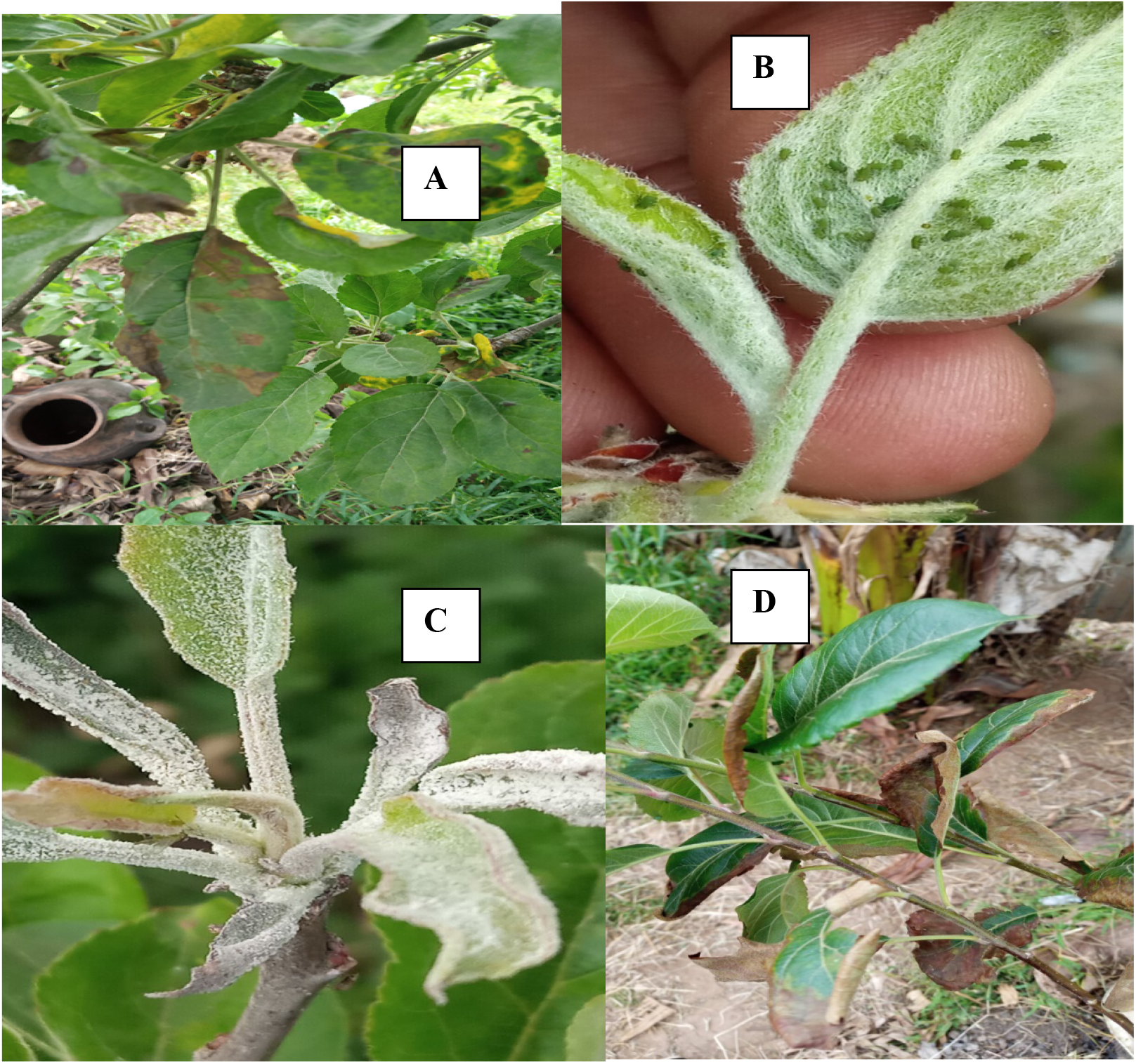
Diseases and insect pest identified in farmers’ apple field. Note; A = Apple scab (*Venturia inaequalis*), B = Green apple aphid (*Aphis pomi*), C = Powdery mildew (*Podosphaera leucotricha*), and D = Fire blight of apple (*Erwinia amylovora*)

The survey of apple fruit tree pests on farmers apple orchards, that carried out from November 2021-April 2021 indicated that, invasion of apple tree leaves by powdery mildew in both surveyed Districts of Silte Zone was significant and severity level was extensive, except Duna Kenema *Kebele* administration of Mirab azernet District and followed by apple scab (with moderate severity level in two Districts), and blight with more slight and moderate severity level. The highest severity percentage of powdery mildew was recorded from Alich wuriro District. During our survey study in the farmers apple orchard, only green apple aphid was recorded from insects pests that infest apple and reported by different researchers in Ethiopia and its infestation on apple leaves was significantly varied from District to District and damage level was again differed from very low to medium. Invasion of apple fruit by apple scab was comparatively high and its severity level was moderate in both surveyed Districts and followed by powdery mildew with varied severity level (slight to moderate). Infestation of insect pests on apple fruit was not recorded in the study area (Table 1 and 2).

**Table 1.**
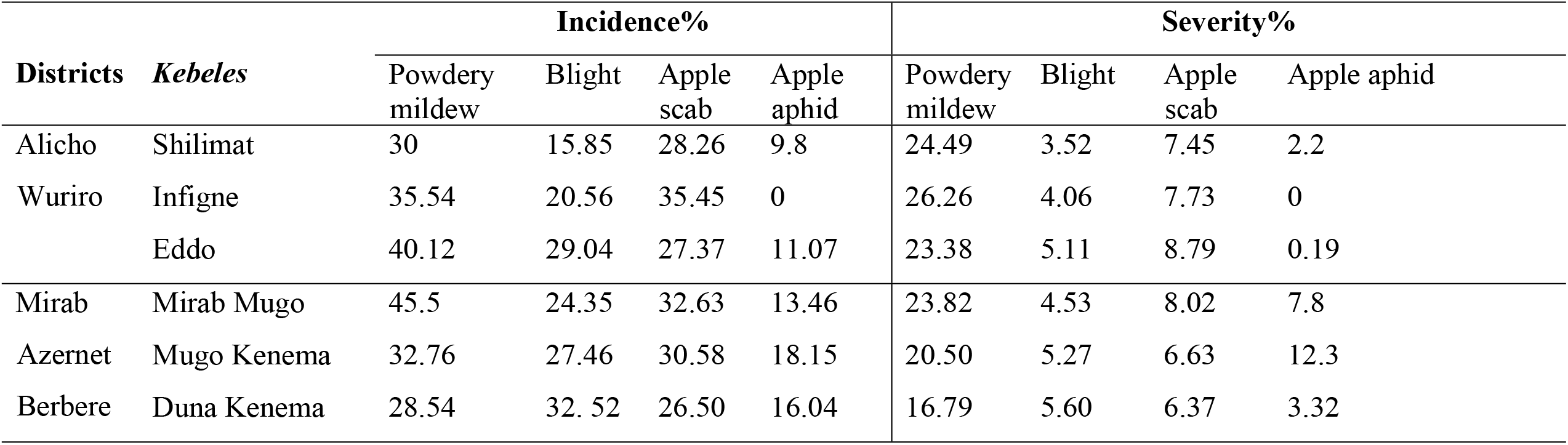
Incidence and severity of diseases and insect pests on apple leaves

**Table 1.** Incidence and severity of diseases and insect pests on apple fruits

| Districts | Kebeles | Incidence% |  |  |  |  |  | Severity% |  |  |  |  |  |
| --- | --- | --- | --- | --- | --- | --- | --- | --- | --- | --- | --- | --- | --- |
|  |  | Powdery mildew | Apple scab | Blight | Maggots | Apple aphids | Caterpillars | Powdery mildew | Apple scab | Blight | Maggots | Apple aphid | Caterpillars |
| Alicho | Shilimat | 15.68 | 20.28 | 0 | 0 | 0 | 0 | 5.24 | 12.52 | 0 | 0 | 0 | 0 |
| Werero | Infigne | 20.32 | 21.64 | 0 | 0 | 0 | 0 | 3.76 | 9.06 | 0 | 0 | 0 | 0 |
|  | Eddo | 18.46 | 26.39 | 0 | 0 | 0 | 0 | 7.48 | 15.11 | 0 | 0 | 0 | 0 |
| Mirab | Mirab Mugo | 23.85 | 24.82 | 0 | 0 | 0 | 0 | 3.82 | 17.53 | 0 | 0 | 0 | 0 |
| Azernet | Mugo Kenema | 12.74 | 21.36 | 0 | 0 | 0 | 0 | 2.50 | 8.27 | 0 | 0 | 0 | 0 |
| Berberere | Duna Kenema | 14.29 | 18.75 | 0 | 0 | 0 | 0 | 6.79 | 5.60 | 0 | 0 | 0 | 0 |

**Table 2.** Educational status of the respondents

| Education level | Percentage of respondents |  |  |
| --- | --- | --- | --- |
|  | Alicho Wuriro | Mirab azernet berber | Over all (zonal) |
| Illiterate | 73.3 | 46.7 | 60 |
| Primary (1-8) | 20 | 40 | 30 |
| Secondary (9-12) | 6.7 | 13.3 | 10 |
| Diploma and above | 0 | 0 | 0 |

### Educational status

As it observed from the result below (Table 3), majority (60%) of apple growers were illiterate in both Districts. Accordingly, it shows there is a great awareness gap between farmers on growing apple fruit tree by using different technologies in the study area. Whereas, from respondents 30% farmers were completed primary school and 10% from respondents were attended secondary school. So, to increase apple fruit tree adoption in the study area; growing apple trees in farmers training center and giving continuous training by agricultural extension workers and none-government organizations is required.

**Table 3.** Growers level of awareness about diseases and insect pests of apple

| Name of District | Which diseases of apple are common in your apple orchard/nursery? |  |  |  |  |  | P-value | Which insect pests of apple are common in your apple orchard/nursery? |  |  |  |  |  |
| --- | --- | --- | --- | --- | --- | --- | --- | --- | --- | --- | --- | --- | --- |
|  |  | Powdery mildew | Apple scab | Blight | Root rot | Total |  | Aphids | Plant bugs | Caterpillars | Not identified | Total | P-value |
| Alichowuriro | N | 25 | 4 | 0 | 1 | 30 | 0.58 | 3 | 0 | 0 | 27 | 30 | 0.041 |
|  | % | 83.3 | 13.3 | 0.0 | 3.3 | 100 |  | 10 | 0.0 | 0.0 | 90 | 100 |  |
| Mierabazernet | N | 18 | 8 | 4 | 0 | 30 |  | 9 | 2 | 0 | 19 | 30 |  |
|  | % | 60 | 26.7 | 13.3 | 0.0 | 100 |  | 30 | 6.7 | 0.0 | 63.3 | 100 |  |
| Over all (%) | N | 43 | 12 | 4 | 1 | 60 |  | 12 | 2 | 0 | 46 | 60 |  |
|  | % | 71.7 | 20 | 6.7 | 1.7 | 100 |  | 20 | 3.3 | 0.0 | 76.7 | 100 |  |

N = Number of respondents

**Table 4.**
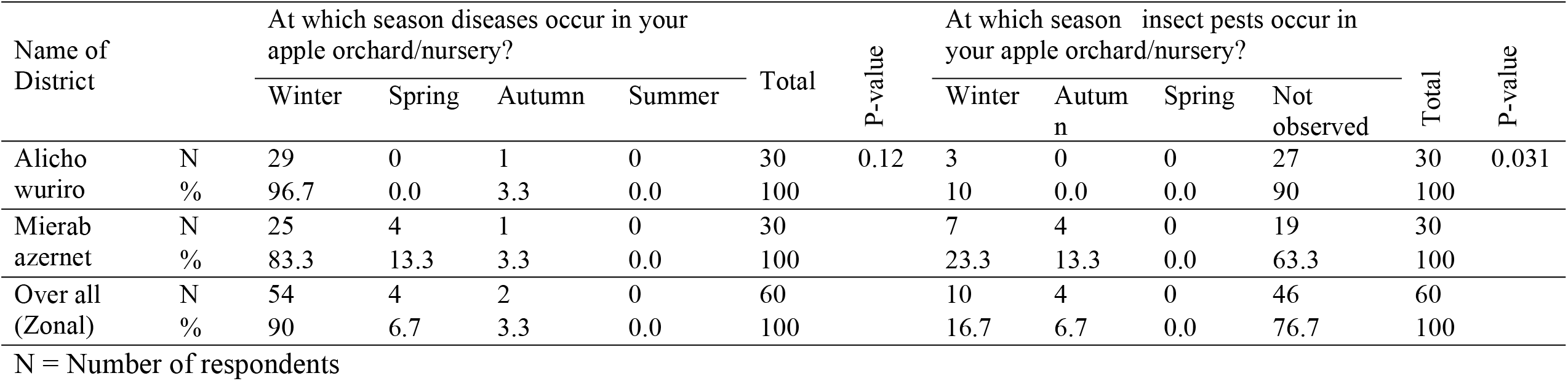
Seasonal distribution of diseases and insect pests in farmers’ apple orchard

### Awareness about diseases and insect pests

Survey output presented below in Table (4) shows that, from respondents in Alicho Wuriro District 83.3% answered powdery mildew as number one disease that affect apple production, followed by 13.3% apple scab and 3.3% root rot. Similarly, in Mierab azernet District from respondents 60% were replied on powdery mildew of apple tree, 26.7% answered apple scab and 13.3% again selected blight of apple and zonally 71.7% respondents were responded powdery mildew as number one disease of apple that hinders apple/pome production in their field, followed by apple scab (20%), blight of apple (6.7%) and root rot (1.7%). Significance difference was not observed between two Districts on occurrence of different apple diseases in their apple orchard (p > 0.05). But, for insect pest in Alicho wuriro only 10% from respondents reacted on aphid and 90% respondents answered they didn’t identified insects that attack their apple fruit tree. However, in Mierab azernet District from respondents 30% reacted on apple aphid, followed by 6.7% plant bugs and also 63.3% from respondents replied that they didn’t observed any insect pest in there apple orchard. Zonally, 76.7% of producers were answered that they didn’t identified insect pests that attack their apple fruit tree and 20% from respondents replied that aphids were attacking their apple tree, followed by 3.3% plant bugs and also significant different was observed between two Districts on occurrence of insect pests (p<0.05).

### Farmers response to seasonal distribution of diseases and insect pests in their apple orchard

The survey result indicated that, significant different was not observed between two Districts on season of disease occurrence (p > 0.05). But, significant different was observed between two Districts on season of insect pest occurrence in farmers apple orchard (p<0.05) (Table 5). Growers from Alicho wuriro District reacted that, disease occurrence in their apple orchard/nursery was observed more on winter season covering (96.7%), followed by autumn (3.3%). In addition to this, in Alicho wuriro about 90% respondents were replied that they didn’t identified insect pests that occur in different seasons and only 10% from the respondents answered that insect pests are occurring in winter season in their apple orchard. But, as per respondents answer it shows that, in Mirab azernet District diseases occur more in winter season and followed by spring and autumn (83.3%, 13.3% and 3.3%) respectively. And similarly, high insect infestation was occur in winter followed by autumn (23.3% and 13.3%) and from respondents 63.3% replied that they didn’t observed/identified any insect pest in their apple orchard in different seasons (Table 5). Zonally, out of the total respondents 90% responded that, winter season was primary period for disease occurrence and similarly 16.7% respondents selected winter season for insect pest infestation in their apple orchard/nursery. As our field observation, from the last week of autumn to the beginning of winter season occurrence of pests were increased due to more leaf proliferation of apple fruit tree and raise of temperature in these seasons and it directs awareness should be provided for growers from agricultural extension workers/other agents in these periods.

**Table 5.** Control measures taken by farmers for pests

| Name of District |  | What control measure do you take for disease? |  |  |  |  | P-value | What control measure do you take for insects? |  |  |  |  | P-value |
| --- | --- | --- | --- | --- | --- | --- | --- | --- | --- | --- | --- | --- | --- |
|  |  | Cultural | Mechanical | No idea | Chemical | Total |  | Cultural | Mechanical | Chemical | No idea | Total |  |
| Alichowuriro | N | 6 | 4 | 20 | 0 | 30 | 0.64 | 2 | 0 | 0 | 28 | 30 | 0.06 |
|  | % | 20 | 13.3 | 66.7 | 0.0 | 100 |  | 6.7 | 0.0 | 0.0 | 93.3 | 100 |  |
| Mierabazernet | N | 2 | 11 | 17 | 0 | 30 |  | 2 | 5 | 0 | 23 | 30 |  |
|  | % | 6.7 | 36.7 | 56.7 | 0.0 | 100 |  | 6.7 | 16.7 | 0.0 | 76.7 | 100 |  |
| Over all (Zonal) | N | 8 | 15 | 37 | 0 | 60 |  | 4 | 5 | 0 | 51 | 60 |  |
|  | % | 13.3 | 25 | 61.7 | 0.0 | 100 |  | 6.7 | 8.3 | 0.0 | 85 | 100 |  |

N = Number of respondents

### Farmers response to diseases and insect pest control measures taken

As presented in Table (6) below, it shows that apple growers were using different management methods to control apple diseases and insect pests in the study area and also significant different was not observed between two Districts on their different management strategies for diseases and insect pests. Zonally, out of the respondents 61.7% and 85% (for diseases and insect pests) responded that they don’t have any idea/information about management methods and in addition to this from the respondents only 25% and 8.3% (for diseases and insect pest) were using mechanical methods like; cutting the infested part of the crop and hand picking to control different pests. However, a small number of respondents were using cultural methods to control diseases and insect pests (13.3% and 6.7%) in the study area.

**Table 6.** Farmers’ response to pruning of apple tree and its season

| Name of District |  | Do you perform pruning of apple? |  |  | P-value | At which season you prune your apple tree? |  |  | Total | P-value |
| --- | --- | --- | --- | --- | --- | --- | --- | --- | --- | --- |
|  |  | Yes | No | Total |  | Summer | Spring | Not applied |  |  |
| Alichowuriro | N | 3 | 27 | 30 | 0.23 | 3 | 0 | 27 | 30 | 0.01 |
|  | % | 10 | 90 | 100 |  | 10 | 0.0 | 90 | 100 |  |
| Mierabazernet | N | 6 | 24 | 30 |  | 6 | 0 | 24 | 30 |  |
|  | % | 20 | 80 | 100 |  | 20 | 0.0 | 80 | 100 |  |
| Over all (Zonal) | N | 9 | 51 | 60 |  | 9 | 0 | 51 | 60 |  |
|  | % | 15 | 85 | 100 |  | 15 | 0.0 | 85 | 100 |  |

N = Number of respondents

### Farmers level of awareness about pruning of apple

During our field observations, except a few apple trees in farmers orchard all are not pruned and also a significant different was not observed between two Districts (p > 0.05) (Table 7). About 85% of respondents answered that, they don’t prune their apple fruit tree in the optimum season and also they don’t know its importance and the method.

## Discussion

As it observed from the result, apple trees in two apple growing Districts were diseased or infested by apple powdery mildew and apple scab from diseases and apple aphid from insects. Similar to our result, two diseases namely, apple scab and powdery mildew were also recorded from North Shewa zone, Amhara region and they caused significant decline on apple production [19]. In line with our result; apple scab, powdery mildew, aphids and caterpillars were recorded from Chencha and Bonke Districts of Gamo Zone and they were infested both leaf and fruit part of apple fruit tree in these Districts [18]. Similar output was reported by [20], and the author directed pests (like; weeds, diseases and insects), lack of information on apple pest control, agronomy, meteorological conditions, and deficient of income to buy seedlings are major constraints of apple fruit tree adoption and production.

Earlier experiences shared by different model farmers in Ethiopia and reported by [21] shows that, growing different crops with apple fruit tree (specially covering the free space by nitrogen fixing crops) is good if growers are educated and care is taken. Similar to this, during our field observation, majority of apple growers in two Districts were intercropped apple fruit trees with different vegetables like; carrot, beetroot, potato, cabbage, garlic and *enset* (*Ensete ventricosum*) and some field crops like; pea, bean and barely without adjusting the space between apple tree and other crops. Similar output is also reported by [23] apple growers in Chencha District intercropped apple fruit tree with other crops. If care is not taken, this type of farming contributes a big share for occurrence of pests and plays a great role by creating suitable environment for pest spreading from one apple tree to the other. Mainly, it creates favorable condition for spreading of powdery mildew and the disease caused weakened/stunting of trees and curling of leaves in the study area. And also the recorded insect pest was more populated on shoot/top part and apical buds of apple trees and caused leaf curling in the study area.

The output of the study agrees with assessment research conducted by [23] and they reported that from apple growers interviewed in Chencha District, Gamo Zone, majorities were not identified insect pests that attack apple fruit trees and also they were not informed about insect pest control measures and only 17.6% from apple growers reacted on green apple aphid. And also the researchers reported powdery mildew and apple scab as number one fungal diseases that familiarized apple fruit tree and identified by growers in this District.

A research conducted by [19] shows that, the number of apple pests increase in hottest meteorological conditions and it has distinctive implication on incidence, severity and seasonal distribution of diseases and insect pests.

Except few growers of apple, majority of growers in surveyed area were not applied the recommended cultural practice (like; inter and intra row between plants, plowing field to destroy alternative hosts, pit preparation, planting garlic on the free space, mulching and etc.) that specified by different experts and model farmers (like; Abiye Astatike, Gurmessa Anissa and Timoteos Hayeso) and reported by [21] for apple fruit tree production. In addition to these, experts recommend uprooting and burning of infected trees if there is severe disease occurrence To control diseases and insect pests of apple, timely pruning plays a great role by interrupting pest movement from one tree to the other. Similar output is reported by [23], most of apple growers in Chencha District don’t prune their apple fruit trees. It indicates, there was information gap between growers and agricultural experts in the study area and apple producers didn’t get enough information about apple fruit tree; improvement, agronomy and protection. During our field observations, powdery mildew was localized on leaves, shoots and on fruits and caused stunted growth of apple trees. On the other hand, aphids were localized on newly sprouting shoots of apple tree and caused curling of leaves and these all were due to not pruning out apple fruit trees timely.

Apple fruit trees are grown in different highland parts of Ethiopia to generate income, improve nutritional status, and enlarge farming system. Silte Zone highland Districts namely; Mirab Azerent and Alicho Wuriro were also producing apple in a great potential from other highland Districts that found in this Zone. Hence, the current assessment research work was conducted with the objective of assessing prevalence and damage of diseases and insect pests of apple and growers experience on pest identification and its management during 2021/21 cropping season in two Districts of Silte Zone. The result of current assessment revealed that, awareness on crop agronomy, pest type and its infestation season, and management measures that carried out by growers in the surveyed area was not satisfying. In both surveyed Districts majority of apple fruit trees were planted under shades of other trees, *enset* (*Ensete ventricosum*) and near to traditional houses except, in those model farmers field [S1 Fig]. During our survey study, powdery mildew, apple scab, blight and green apple aphid were recorded diseases and insect pest of apple fruit tree. From these, powdery mildew and apple scab were more damaging diseases of apple fruit tree and they imply economic threat if not managed and also green apple aphid was the only recorded insect pest that caused leaf curling of apple fruit tree in the surveyed area. About 71.7% of respondents were responded powdery mildew as number one disease of apple that hinders apple production in their field. About 76.7% of producers were answered that they didn’t identified insect pests that attack their apple fruit tree. Out of the total respondents 90% responded that, winter season was primary period for disease occurrence and similarly 16.7% respondents selected winter season for insect pest infestation in their apple orchard/nursery. Out of the respondents 61.7% and 85% (for diseases and insect pest) responded that they don’t have any idea/information about management methods. About 85% of respondents answered that, they don’t prune their apple fruit tree in the optimum season and also they don’t know its importance and the method. Based on the results of our survey, we recommend the followings; apple trees should be planted in farmers training center and continuous training about tree improvement, agronomy and protection should be given by agricultural extension workers for producers, growers should be informed about break of apple tree dormancy, sprouting, flowering, fruit set, and maturity and also time of pest infestation and control measures, in all apple fields surveyed newly emerging shoots of apple were more infested by pests, therefore early cutting of infested shoot/leaf is recommended to stop further dispersion of these pests, apple plantation should be encouraged in fields free from any other crop and more extensive studies are advised to exhaustively determine the biology and management of major pests that attack apple fruit tree to manage the pests effectively, efficiently and economically.

## Supporting information

**S1 Fig. Apple fruit tree intercropped with other crops**

**S1 Table. Questionnaire**

## Acknowledgements

At the beginning we express sincere thanks to our Almighty God. Our genuine thanks go to workers of Mirab azernet and Alicho wuriro District Horticulture experts. We would like also to thank the development agents of each *Kebele* administrate for their kindness and patience in providing the relevant information during the study.

## Author contributions

Identifying the thematic area: Sebsibe Sh. Woto. Identifying suitable methodology to justify research hypothesis and organizing information from different sources: Sebsibe Sh. Woto and Adimasu A. Aloto. Undertaking the analysis and wrote the paper: Sebsibe Sh. Woto and Adimasu A. Aloto.

